# Analyzing the relationship between distracted driving and eye movement using multimodal data collected during car driving

**DOI:** 10.1101/2021.12.31.474674

**Authors:** Daigo Uraki, Kensuke Tanioka, Satoru Hiwa, Hiroshi Furutani, Tomoyuki Hiroyasu

**Affiliations:** Graduate school of Life and Medical Sciences, Doshisha University, Japan; Department of Biomedical Sciences and Informatics, Doshisha University, Japan; AI x Humanity Research Center, Doshisha University, Japan

**Keywords:** driver distraction, driving simulator, multimodal dataset, gaze, steering angle, perinasal perspiration

## Abstract

Human error is the leading cause of traffic accidents and originates from the distraction caused by various factors, such as the driver’s physical condition and mental state. One of the significant factors causing driver distraction is the presence of stress. In a previous study, multiple stressors were used to examine distraction while driving. Multiple stressors were given to the driver and the corresponding driver biometric data were obtained, and a multimodal dataset was published thereafter. In this study, we reiterate the results of existing studies and investigated the relationship between gaze variability while driving and stressor intervention, which has not yet been examined. We also examined whether biometric and vehicle information can estimate the presence or absence of secondary tasks during driving.

## 1 Introduction

In today’s society, automobiles are an indispensable part of our lives. However, they are also a factor in many traffic accidents. One of the causes of traffic accidents is human error due to distraction caused by various factors, such as the driver’s physical condition, mental state, and external environment. Distraction is a state in which resources are not fully invested in the task at hand. Humans usually cope with tasks through moderate sympathetic responses. When stress is added to the original task, it excites the sympathetic nervous system, depriving the original task of the resources it needs. This lack of resources leads to distractions.

Stressful stimuli are referred to as stressors. Stimuli while driving are sensory-motor stressors, with stressors resulting from operating devices, such as texting while driving being the most typical [1–3]. In addition to sensory-motor stressors, cognitive and affective stressors have also been reported [4]. To avoid stress-induced distraction, it is necessary to recognize and eliminate the presence of stressors. In this study, instead of evaluating whether a stimuli is a stressor by observing external factors, we investigated the presence of a stressor using biometric and vehicle information of the driver while driving. If a stressor can be detected by obtaining biometric and vehicle information, the development of a system that alerts the driver, becomes possible. Skin sweating, pupil diameter, and brain activity information, such as changes in the cerebral blood flow and electroencephalograph, which were measured by functional near-infrared spectroscopy (fNIRS) and electroencephalogram, have been used as biometric information for this purpose.

“A multimodal dataset for various forms of distracted driving” is a dataset in which biometric and vehicle information obtained from controlled experiments using a driving simulator is measured [5]. This dataset contains the experimental results of 68 people who drove the same highway simulation under 4 different conditions with given stressors (cognitive, emotional, and sensory-motor). This previous study confirmed that cognitive, emotional, and sensory-motor stressors caused a significant increase in each of the mean sympathetic nervous systems that the perinasal electro dermal activity(EDA) could examine. It was also confirmed that the stressors caused a significant increase in the mean absolute steering volume [6].

The perinasal EDA is closely related to the parasympathetic nerve response. In contrast, in addition to the perinasal EDA, this multimodal dataset contains pupil diameter information related to the parasympathetic response, data that tracks the coordinates in the space of the gaze during driving, and palm EDA. In a previous study, biometric data were measured with multiple modalities, but only perinasal EDA data were used. It is also interesting to note the influence of measured data other than perinasal EDA data and stressors on driving. In some cases, it is assumed that perinasal EDA cannot be acquired correctly because of the influence of body hair, and other biometric data may be easier to measure in an actual driving environment.

Therefore, we re-analyzed the data used in existing studies and conducted an additional study using ocular information, which is easy to measure in actual driving environments. In summary, we hypothesized as follows: 1) Gaze variability during driving increases with stressor intervention, and 2) The presence of secondary tasks during driving can be discriminated using biometric and vehicle information.

## 2 Materials and Methods

This study uses “A multimodal dataset for various forms of distracted driving”. The details of the data used are described in this section.

### 2.1 Participants

Participants were those who held a driver’s license, had been driving for more than 1.5 years, and had a standard or corrected vision. Participants were divided into two groups: a younger group (18–27 years old) and an older group (60 years old and above). Data were obtained from 68 participants (35 men/33 women), excluding those not correctly recorded. Perinasal sweating data were not obtained for 9 of the 68 participants. In this study, participants for whom all data could be measured were selected. Therefore, participants were selected from the total dataset when all the data could be measured. As a result, there were 37 participants (13 men/24 women).

The experimental design consisted of two designs. The designs for Experiments 1 and 2 are presented below.

### 2.2 Design of Experiment 1

In Experiment 1, the participants were asked to perform driving tasks under six conditions. The Cognitive Drive(CD), Emotional Drive(ED), and Sensory-motor Drive(MD) tasks were conditions in which a secondary task was given while the Neutral Drive(ND) task was a condition in which no secondary task was given during driving. After the Practice Drive(PD) and Relaxing Drive(RD) tasks were performed, five phases were conducted. The phases during the implementation of the driving task LDj(j∈[C,E,M]) under stress conditions are as follows:

- Phase P1: Driving without distraction for about 80 s
- Phase P2: Driving while performing a secondary task for about 160 s
- Phase P3: Driving without distraction for about 240 s
- Phase P4: Driving while performing the secondary task for about 160 s
- Phase P5: Driving for approximately 120 s without the diversion of attention

The order of the secondary tasks was determined randomly and performed by each participant. The details of the driving task performed by the subject are as follows.

#### 2.2.1 Practice Drive(PD) Task

The participants drove on an 8 km straight section of a 4-lane highway at the prescribed speed limit. Two lanes each way, and the participant’s car drove in the right lane (R). The speed limit changed every few kilometers (80 km/h→50 km/h→100 km/h).

#### 2.2.2 Relaxing Drive(RD) Task

In the RD task, the subject drove a 10.9 km straight section of a four-lane highway at a speed limit of 70 km/h. There were two lanes in each direction, and the participants drove in the R lane. The driving conditions were set such that the oncoming lane would not be congested (maximum three vehicles/km). After 5.2 km of driving, the participants were allowed to change lanes from right to left. After driving at least 1.2 km in the left lane, those who wished were allowed to return to the right lane. This lane change was incorporated into the experiment to reduce the monotony of driving.

#### 2.2.3 Neutral Drive(LD_*ϕ*_=ND) Task

The participants were forced to drive with a non-driving load during the LD task. The load was given as a secondary task during the driving. The ND task was performed under driving conditions when there was no load. The following driving condition is called the load driving (LD) condition: The participants drove on a 10.9 km four-lane highway with a speed limit of 70 km/h. Two lanes were dedicated to each traffic, and the subject drove in the R lane. Traffic congestion in the oncoming lane was more than 12 vehicles/km. The left lane was under construction, and the R lane was provided with traffic signs. At the 4.4-km point, the participants were forced to change from the R lane to the left lane, and after driving in the left lane for 1.2 km, they returned to the R lane. During this detour, a construction cone appeared on the right side of the lane.

#### 2.2.4 Cognitive Drive(LD_*C*_=CD) Task

In addition to the ND condition, the CD task involved driving under cognitive stress. The cognitive stressor was divided into two phases: mathematical and analytical. In each phase, the experimenter started the questions verbally at the beginning of the list. Finally, the experimenter stopped the questions. Participants were required to answer all the questions to the best of their abilities. The mathematical and analytical questioning phases were randomized and administered to the participants.

#### 2.2.5 Emotional Drive(LD_*E*_=ED) Task

In addition to the ND condition, the ED task involved driving under emotional stress. Emotional stressors were divided into two levels. The experimenter verbally asked for an emotional stressor. There were two sets of questions: one sharp and one relatively less sharp. During the first stressful phase, basic questions topping the relevant list were asked for 20 s. For the remainder of the first stressful phase, the experimenter asked a sharp question from the relevant list. In the second stress phase, the basic questions were asked for 30 s from the questions left over from the first stress phase. For the remainder of the time, sharp questions were asked from the same list of questions left over. Participants were asked to answer all of these questions to the best of their ability.

#### 2.2.6 Sensorimotor Drive(LD_*M*_ =MD) Task

In the MD tasks, driving was performed under sensory-motor stress, in addition to the ND condition. Participants replied to the text messages received on a smartphone.

### 2.3 Design of Experiment 2

Following Experiment 1, Experiment 2 was conducted using the same driving simulator. In Experiment 2, a failure drive (FD) task was conducted. The details of the FD task are as follows:

At the end of Experiment 1, the participants drove 3.2 km on a highway. The participants were divided into two groups: a group without a secondary task (y = o) and a group driving the last 2 km under mixed stress with a secondary task (y = L).

The participants in y = L first typed the text displayed on their smartphones. Subsequently, while the participants were texting, the experimenter verbally asked the mathematical, analytical, and emotional questions to which the participants responded. At the end of the driving session, all participants stopped at a red light at an intersection.

In this task, unintended acceleration occurred before the light turned green, causing the vehicle to collide with another vehicle entering the intersection. Before the collision, the reaction time was five seconds.

A summary of the three sessions included in the FD is as follows:

- Phase P1: First session of driving - no distraction
- Phase P2: After 1.2 km – y = o no distraction; y = L mixed distraction
- Phase P3: Unintended acceleration event for about 11 s

## 3 Analysis Methods

To investigate the effects of the secondary task, the following analysis was conducted. In each running phase of the LDj(j∈[C,E,M]) task under each stress condition subjected to the secondary task, the index value of each phase was calculated using the following equations (1 to3):

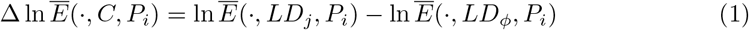

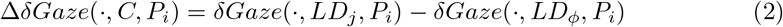

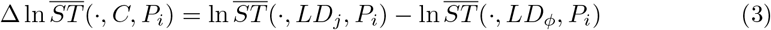

For each of the five phases of ED, CD, and MD, a one-way ANOVA was performed on the calculated values of the five phases. A two-sample t-test with multiple comparison corrections will be conducted between each phase if a statistically significant difference is found. For the multiple comparison correction, the Bonferroni method was used.

### 3.1 Estimation of the presence of secondary issues

The FD task was used to classify participants with and without secondary tasks based on their attributes, vehicle information, and biometric data. In the classification, gender and age, which have been suggested to influence driving maneuvers in previous studies, were used as participant attributes. The absolute value of the steering angle during the Phase 2 section of the FD task, where the group with the sub-task engaged in driving under mixed stress, was used. In addition, perinasal EDA, which is strongly related to the parasympathetic nervous system, and horizontal and vertical gaze dispersion were used as biometric indicators of the participants’ state while driving. These participant attributes, vehicle information, and biometric information were used as explanatory variables, and the presence or absence of the participant’s secondary task was used as the objective variable. Since the objective variables were discrete binary values, logistic regression was used to conduct the principal regression. The regression equation is given by Equation (4).

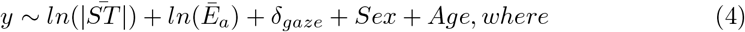

- ln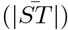: Natural logarithm of the mean value of the absolute steering angle adjusted to baseline at rest for each subject
- ln(*Ē*_*a*_): Natural logarithm of mean perinatal EDA value adjusted to baseline at rest for each subject
- *δ*_*gaze*_: Variance of the horizontal or vertical gaze coordinates of each subject
- Sex: biological sex (male vs female)
- Age: Age group (Young: < 27 vs Old: > 60)

The leave-one-out cross-validation was also used to evaluate the logistic regression results. We called it true positive (TP) when the correct answer was loaded and the predicted value was loaded; true negative (TN) when the correct answer was not loaded and the predicted value was not loaded; false positive (FP) when the correct answer was not loaded and the predicted value was loaded; and false negative (FN) was defined when the correct answer was loaded and the predicted value was not loaded. The evaluation items are accuracy (Equation (5)), precision (Equation (6)), detection rate (Equation (7)), and F-value (Equation (8)).

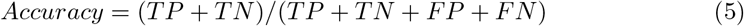

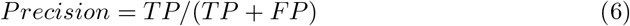

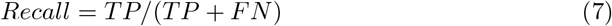

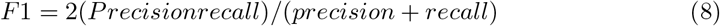

### 3.2 Estimating Steering Angle

Two regression models were used to estimate the mean absolute value of the steering angle in the FD task.

### 3.3 Model 1

Predictions were made based on the perinasal EDA, age of the participants, and presence or absence of a secondary task load. This prediction is a reiteration of the results of previous studies. The regression equation for the steering angle prediction is given by Equation (9).

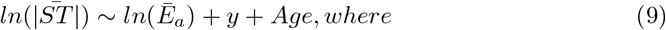

- ln(*Ē*_*a*_): Natural logarithm of mean perinatal EDA volume adjusted to baseline at rest for each subject
- y: Load on secondary tasks(y= o vs. y=L)
- Age: Age group(Young: < 27 vs Old: > 60)

### 3.4 Model 2

Model 2 is a model that uses horizontal and vertical gaze information as biometric information instead of perinasal EDA, which was the focus of the previous study. The regression equation for Model 2 is shown in Equation (10).

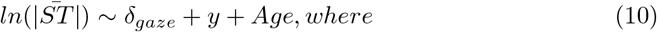

- *δ*_*gaze*_: Variance of the horizontal or vertical gaze coordinates of each subject
- y: Load on secondary tasks(y= o vs. y=L)
- Age: Age group(Young: < 27 vs Old: > 60)

## 4 Results

### 4.1 Analysis on Loaded Drivej(j∈[C,E,M]) Data

The effects of the presence or absence of the secondary task in Experiment 1, with the results of the CD, ED, and MD tasks are shown in Figure 2, 3, and 4, respectively.

**Figure 1.**
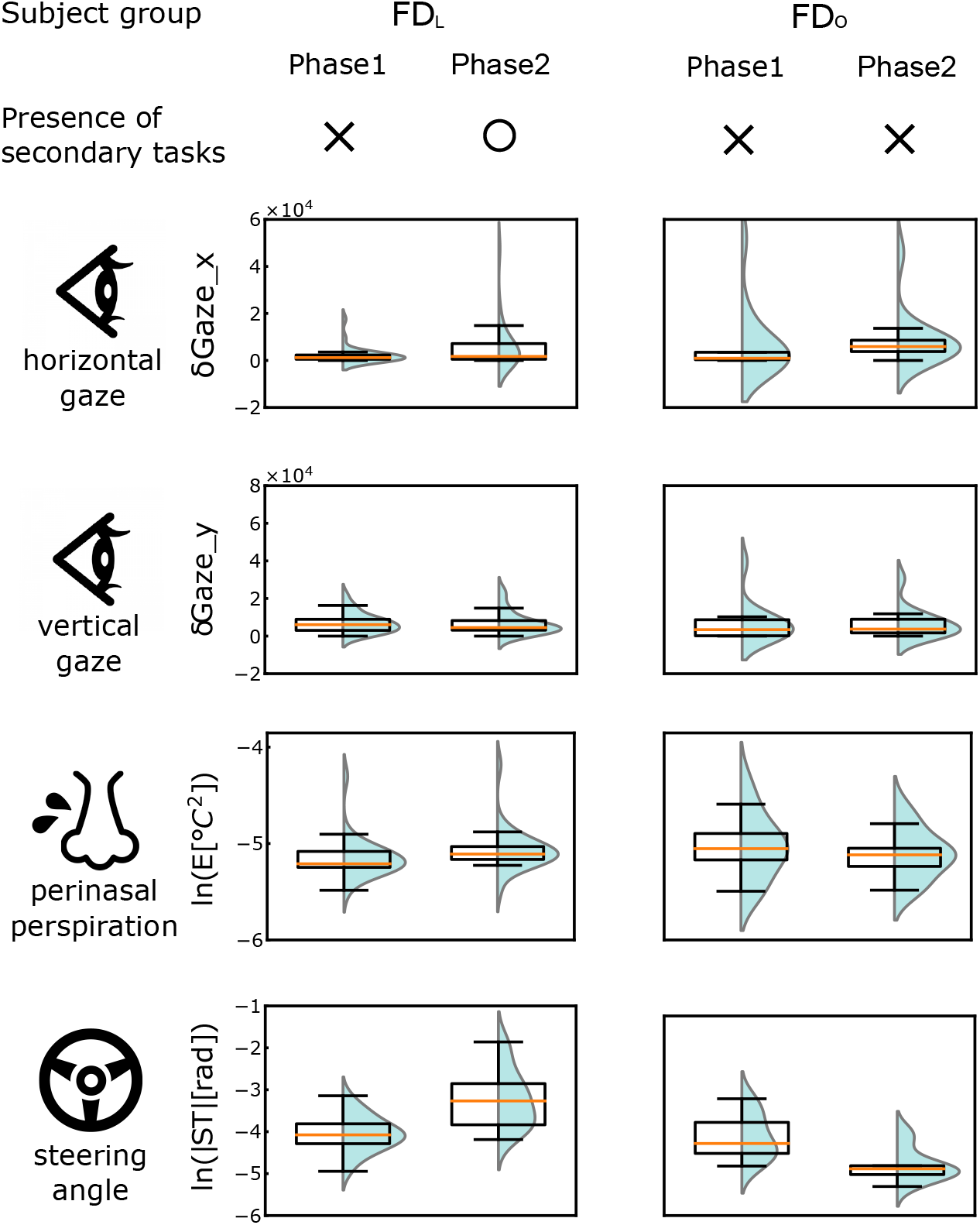
Biometric and vehicle information for each phase during FD task

**Figure 2.**
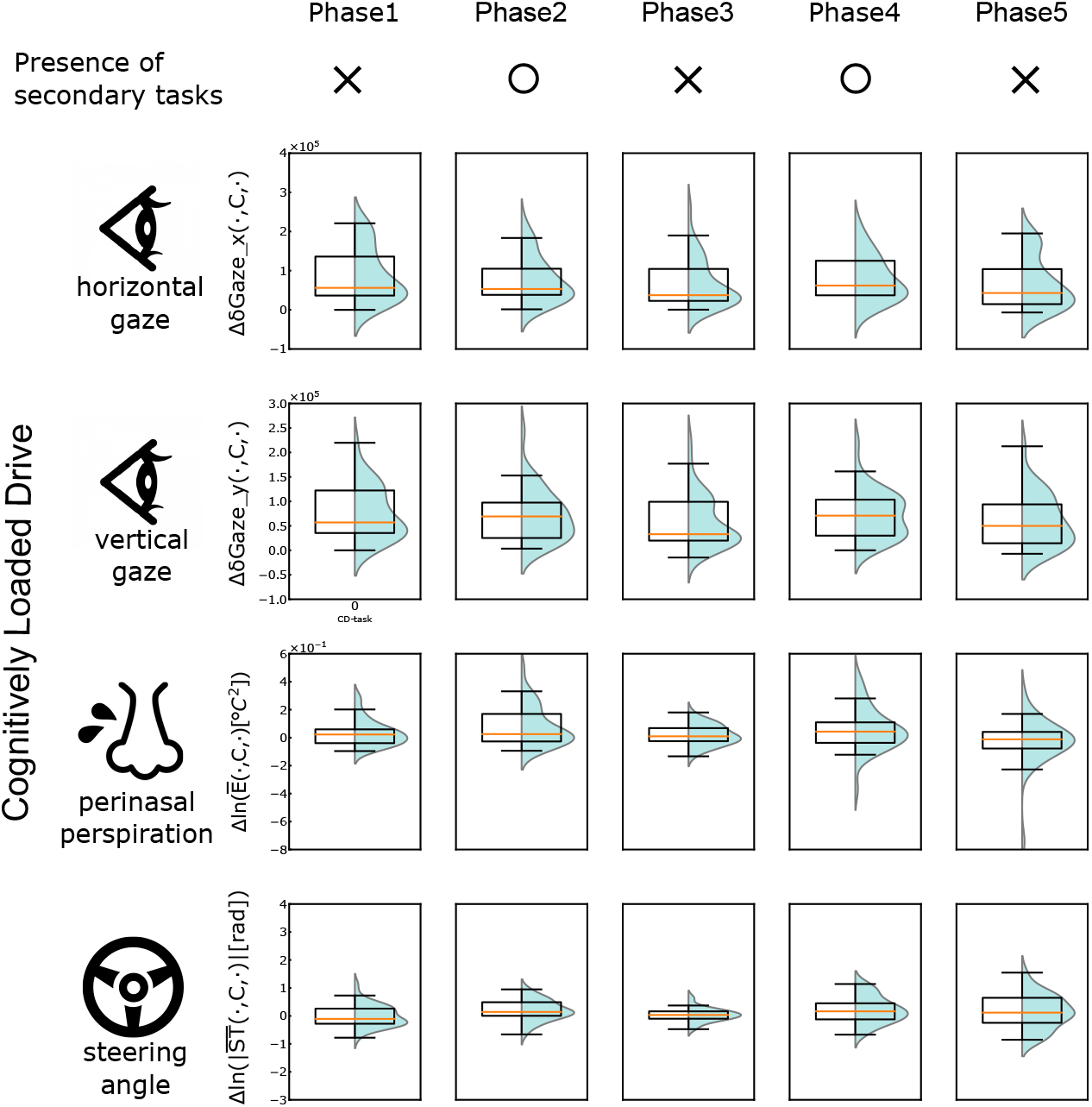
Biometric and vehicle information for each phase during CD task

**Figure 3.**
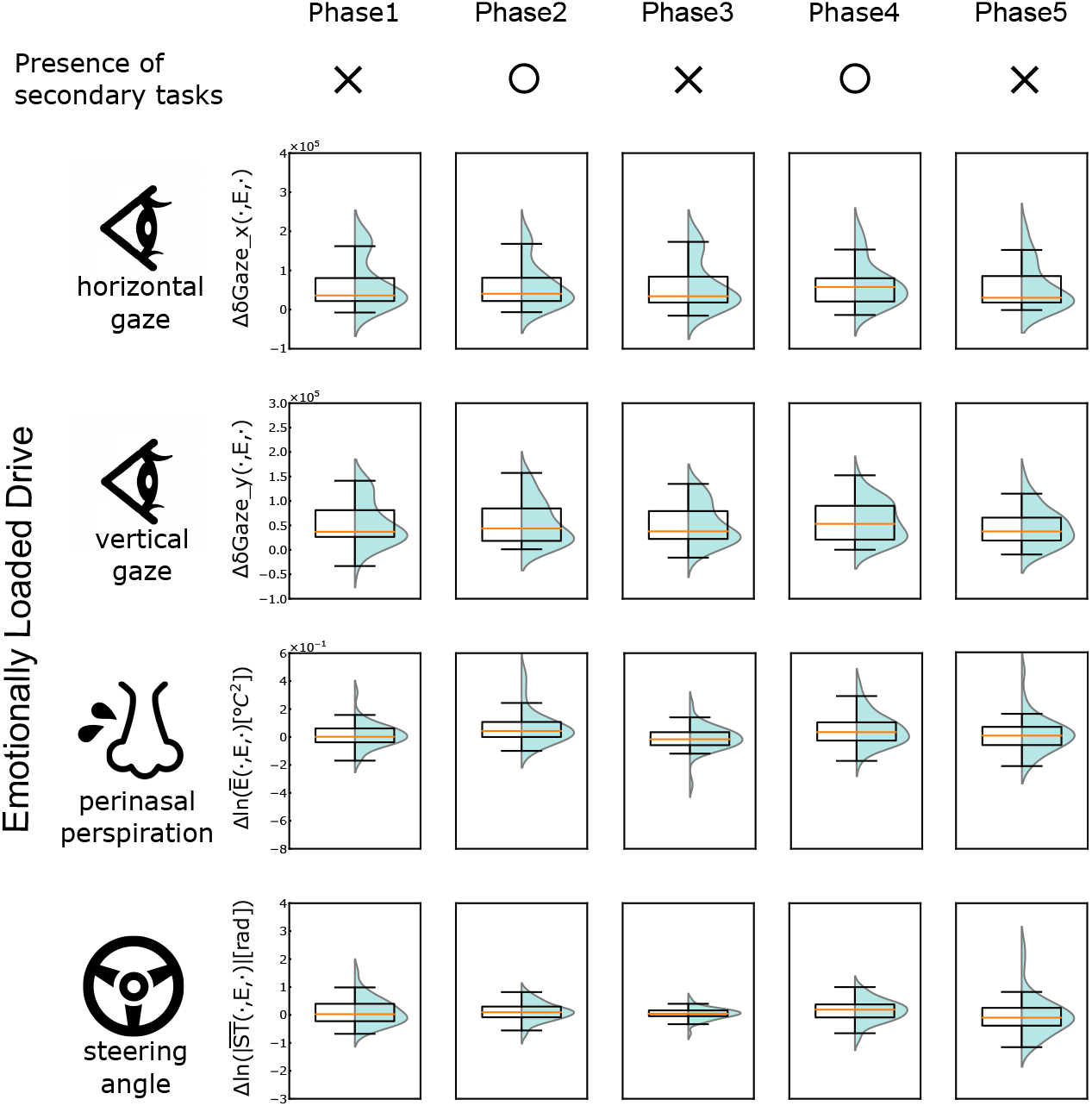
Biometric and vehicle information for each phase during ED task

**Figure 4.**
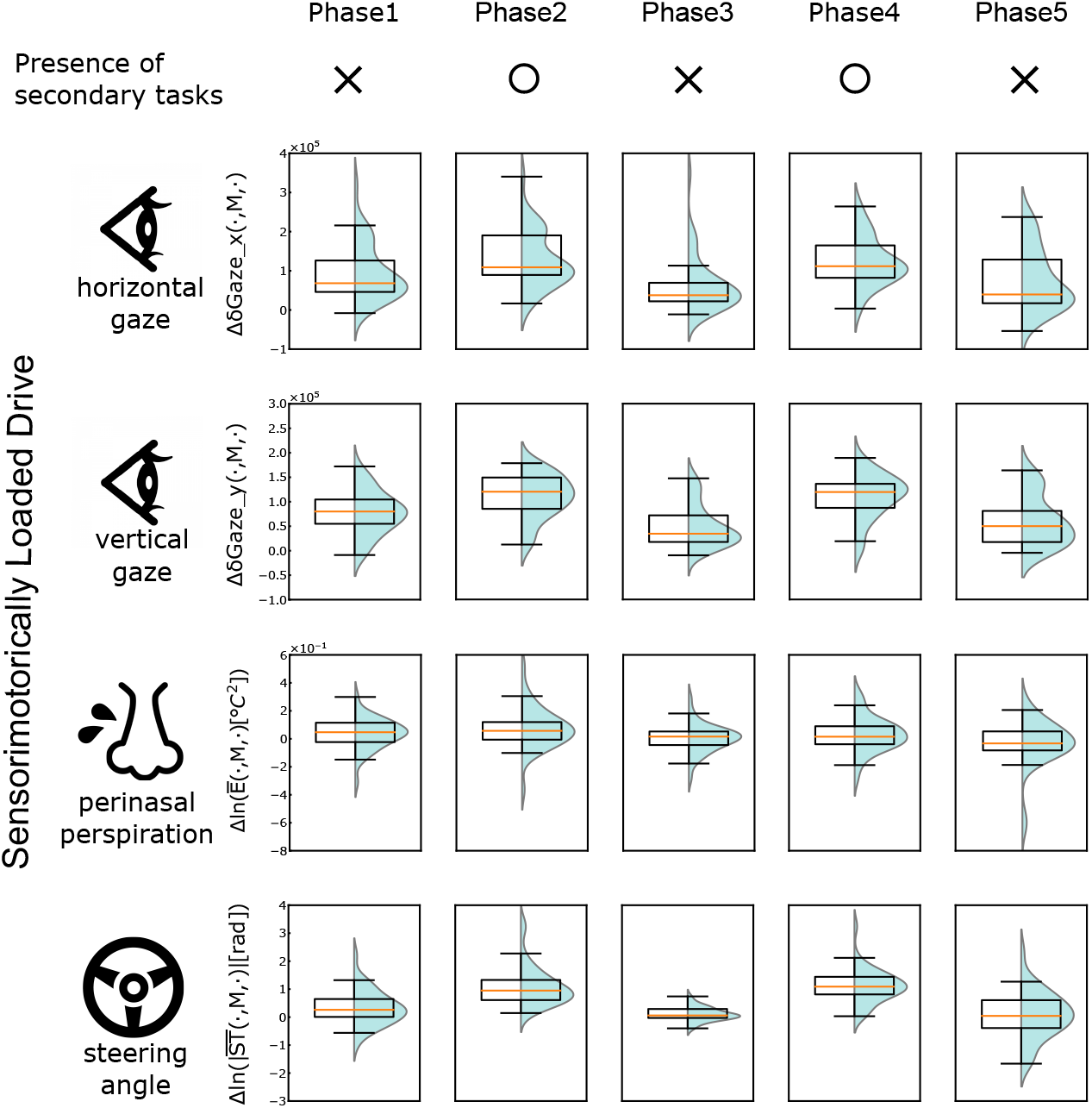
Biometric and vehicle information for each phase during MD task

When comparing the phases of the CD and ED tasks with the phases of the ND task, we observed that there is a statistically significant difference in the steering angle in the phases where the subtasks are present (Phase 2 and Phase 4) (CD: *p*<0.01, ED: *p*<0.05). This result is similar to those of previous studies. When comparing each phase of the MD task with each phase of the ND task, there was a statistically significant difference in steering angle in the phases where the sub-task existed (Phase 2 and Phase 4) (*p*<0.001). In Phase 1, there was also a statistically significant difference in steering angle (*p*<0.01). In the previous study, there was a statistically significant difference in steering angle in the phases where the MD task existed (Phase 2 and Phase 4) and in the phases immediately after the MD task (Phase 3 and Phase 5). However, the results of this analysis are not available.

There was a statistically significant difference in the perinasal EDA between Phase 2 of the CD task, Phase 2 and Phase 4 of the ED task, and the section of Phase 2 of the MD task for each phase of the ND task (*p*<0.01). Since all phases in which an increase in perinasal EDA was observed were in the section where the secondary task was performed, it can be said that the results of perinasal EDA were similar to the results of previous studies.

In the phase where the secondary tasks of the CD and ED tasks were performed, there was no significant difference in gaze variance. On the other hand, there was a statistically significant difference in gaze dispersion between the phase in which the secondary task of the MD task was performed and the phase immediately following it (*p*<0.01). This result was confirmed for both the horizontal and vertical components of gaze.

### 4.2 Analysis on Failure Drive Data

#### 4.2.1 Effects of Combined stressors

The biometric and vehicle information for each phase of the FD task is shown in Figure 1. In Figure 1, two groups of analysis results are shown, and divided based on the presence or absence of secondary tasks.

The results of the analysis of the FD task data for the two groups divided by the presence or absence of a secondary task are shown in Figure 1.

In the FD task, there were no significant differences in the perinasal EDA, steering angle, or gaze between the groups divided by the presence or absence of the secondary task.

There was a statistically significant difference in the perinasal EDA and steering angle between Phase 1, in which the participants drove without the secondary task, and Phase 2, in which they drove while performing the secondary task with compound stress (*p*<0.01). However, a statistically significant difference (*p*<0.01) was also found when comparing Phase 1 and Phase 2 in the group without the secondary task. This cannot be ruled out as an effect of secondary tasks.

#### 4.2.2 Estimating the presence of secondary issues

Participants were grouped according to the presence or absence of a secondary task. The group to which the participants belonged was regressed. The confusion matrix of the regression results is presented in Table 1.

**Table 1.** Confusion matrix of regression results

|  |  | True diagnosis |  | Total |
| --- | --- | --- | --- | --- |
|  |  | Positive | Negative |  |
| Screening test | Positive | 22 | 2 | 24 |
|  | Negative | 2 | 11 | 13 |
| Total |  | 24 | 13 | 37 |

The results confirm that it is possible to discriminate the presence or absence of mixed stress of the driver based on the attributes of the driver, such as gender and age, the average steering angle while driving, and the biometric information of the participants (perinasal EDA, eye gaze).

The values of various evaluation items are shown in Table 2.

**Table 2.** Results of logistic regression

|  |  |
| --- | --- |
| Accuracy | 0.891 |
| Precision | 0.846 |
| Recall | 0.846 |
| F1 | 0.846 |

As for the results of logistic regression, it was possible to discriminate the presence or absence of mixed stress on the driver in the driving compartment based on the attributes of the driver such as gender and age, the average steering angle during driving, and the biometric information of the subject (33 out of 37 subjects correctly identified the presence or absence of stress). The values of the various evaluation items were calculated. The values of the various evaluation items are shown in Table 2.

#### 4.2.3 Estimation of Steering Angle

The coefficients of determination obtained when regressing the steering angle are listed in Table 3.

**Table 3.** Prediction results for steering angle

| Biometric data used for regressors | R2 |
| --- | --- |
| perinasal perspiration | 0.716 |
| horizontal gaze | 0.725 |
| vertical gaze | 0.712 |

The coefficients of determination for perinasal sweating, horizontal gaze, and vertical gaze were almost the same. Therefore, there was no difference in the degree of influence between the perinasal EDA and gaze. When the correlation of each regressor was examined, the only regressors that correlated with the prediction of the steering angle were the presence of secondary task load and age (*p*<0.05). On the other hand, perinasal EDA, a biological indicator of the degree of arousal that is thought to be closely related to changes in parasympathetic nerve activity due to the effects of secondary tasks in the experiment, showed no correlation.

In a previous study, perinasal EDA was correlated with mixed stress load and age in the estimation of the steering angle. In contrast, the presence of sensory-motor stress load and age were correlated in the prediction of the steering angle in the present study. In addition, the number of participants in this study was smaller than that in the previous study. Therefore, it should be noted that it is difficult to obtain significant results compared to previous studies.

When regressed using Model 2, the presence of mixed stress load and age was correlated with the steering angle estimation (*p*<0.05). In Model 2, the gaze is used as biometric information as an alternative to perinasal EDA in predicting the steering angle. In other words, it was suggested that age might affect the steering maneuvers.

If data on the presence or absence of mixed stress load and age are obtained, the steering angle can be predicted without the need for biometric information.

## 5 Conclusion

This study indicates that steering maneuvers and perinasal EDA are affected by secondary task interventions, regardless of the type of stress. This result is in line with the findings of previous studies. Gaze variability is also affected by secondary tasks that cause sensory-motor stress. This result supports the hypothesis that gaze variability during driving is increased by stressor intervention when the type of stress is sensory-motor in nature.

The logistic regression results show that when attributes such as the participant’s age and gender are known, it is possible to determine the presence or absence of secondary tasks that cause compound stress while driving using biometric and vehicle data. This result supports the second hypothesis that the presence or absence of secondary tasks while driving can be determined using biometric and vehicle data. However, since the data used to determine the presence or absence of secondary tasks were measured during an experiment using a secondary task that induced combined stress, the same level of results may not be obtained if the presence or absence of a single sensory-motor stress, cognitive stress, or emotional stress-inducing task was used as the objective variable. In addition, if only eye gaze data are used to estimate the driver’s state, it may be challenging to detect cognitive and emotional stressors due to conversations with passengers. Therefore, it is necessary to estimate the driver’s state using vehicle data, such as steering wheel operation and other biometric data.

## Notes

### Competing Interest Statement

The authors have declared no competing interest.

